# Cold stress induces a rapid redistribution of the antagonistic marks H3K4me3 and H3K27me3 in *Arabidopsis thaliana*

**DOI:** 10.1101/2024.02.29.582735

**Authors:** Léa Faivre, Nathalie-Francesca Kinscher, Ana Belén Kuhlmann, Xiaocai Xu, Kerstin Kaufmann, Daniel Schubert

**Affiliations:** Epigenetics of Plants, Freie Universität Berlin, Berlin, Germany; Department for Plant Cell and Molecular Biology, Institute for Biology, Humboldt-Universität zu Berlin, Berlin, Germany

## Abstract

When exposed to low temperatures, plants undergo a drastic reprogramming of their transcriptome in order to adapt to their new environmental conditions, which primes them for potential freezing temperatures. While the involvement of transcription factors in this process, termed cold acclimation, has been deeply investigated, the potential contribution of chromatin regulation remains largely unclear. A large proportion of cold-inducible genes carries the repressive mark histone 3 lysine 27 trimethylation (H3K27me3), which has been hypothesized as maintaining them in a silenced state in the absence of stress, but which would need to be removed or counteracted upon stress perception. However, the fate of H3K27me3 during cold exposure has not been studied genome-wide. In this study, we offer an epigenome profiling of H3K27me3 and its antagonistic active mark H3K4me3 during short-term cold exposure. Both chromatin marks undergo rapid redistribution upon cold exposure, however, the gene sets undergoing H3K4me3 or H3K27me3 differential methylation are distinct, refuting the simplistic idea that gene activation relies on a switch from an H3K27me3 repressed chromatin to an active form enriched in H3K4me3. Coupling the ChIP-seq experiments with transcriptome profiling reveals that differential histone methylation correlates with changes in expression. Interestingly, only a subset of cold-regulated genes lose H3K27me3 during their induction, indicating that H3K27me3 is not an obstacle to transcriptional activation. In the H3K27me3 methyltransferase *curly leaf (clf)* mutant, many cold regulated genes display reduced H3K27me3 levels but their transcriptional activity is not altered prior or during a cold exposure, suggesting that H3K27me3 may serve a more intricate role in the cold response than simply repressing the cold-inducible genes in naïve conditions.

## 1 Introduction

Low temperatures negatively affect both plant growth and productivity. Low temperature stress can be divided into chilling stress (0-15°C for temperate plants such as *Arabidopsis thaliana*) and freezing stress (subzero temperatures) and plants devised strategies to cope with both of these stress types (Zarka *et al*., 2003). While plants have a constitutive tolerance towards chilling stress, the freezing tolerance of most plants growing in a temperate climate is increased during an exposure to low but non-freezing temperatures, a process known as cold acclimation (Gilmour, Hajela and Thomashow, 1988; Jan, Andrabi and others, 2009). Cold acclimation relies on the production of a variety of proteins whose function is to limit the damage caused by a putative future freezing event and is therefore associated with a significant transcriptional reprogrammation (Calixto *et al*., 2018; Shi, Ding and Yang, 2018). Upon perception of low temperature, the ICE1 transcription factor is activated, thereby inducing the expression of the C-repeat Binding Factors (CBFs) (Wang *et al*., 2017). In turn, the CBFs bind to the C-Repeat motifs of cold-responsive (*COR*) genes (Yamaguchi-Shinozaki and Shinozaki, 1994; Medina *et al*., 1999). This results in the transcriptional activation of thousands of *COR* genes within a few hours of exposure to low temperatures. While numerous transcription regulators have been identified as playing a role in cold acclimation, the putative contribution of the chromatin status to this transcriptional reprogramming remains underinvestigated.

Chromatin is an important contributor to the regulation of transcription, as it controls the accessibility of the underlying DNA to the transcriptional machinery. Within the nucleus, DNA is wrapped around octamers of histones, forming the nucleosome, which is the basic organizational unit of the chromatin (Kornberg, 1977; Luger *et al*., 1997). Histones tails protrude from the nucleosome and can be heavily post-translationally modified by acetylation, methylation and phosphorylation, among others (Luger and Richmond, 1998; Zhao and Garcia, 2015). Those histone post-translational marks (PTMs) can affect the transcriptional activity of the underlying gene directly, by modulating the strength of the interaction between DNA and histones, or indirectly, by recruiting other proteins called histone readers that recognize and bind to specific histone PTMs (Blakey and Litt, 2015). Depending on whether they are associated with transcribed or silenced genes, histone PTMs are classified as active or repressive marks, respectively. Some of the most characterized histone PTMs are the trimethylation on lysine 4 (H3K4me3) and 27 (H3K27me3) of histone 3, which respectively act as an active and a repressive mark (Roudier *et al*., 2011; Cheng *et al*., 2020). H3K27me3 is deposited by the Polycomb Repressive Complex 2 (PRC2) and contributes to the silencing of its targets (Müller *et al*., 2002; Zhang *et al*., 2007). PRC2, which was initially identified in *Drosophila*, consists of four subunits, including the Enhancer of zeste (E(z)) methyltransferase (Müller *et al*., 2002). Three homologs of E(z) have been identified in *Arabidopsis thaliana*: CURLY LEAF (CLF), SWINGER (SWN) and MEDEA (MEA) (Chanvivattana *et al*., 2004). The action of PRC2 is counteracted by methyltransferases from the Trithorax (TrxG) group, which deposit H3K4me3 (Ingham, 1983; Ringrose and Paro, 2004). H3K27me3 and H3K4me3 have long been described as being mutually exclusive, with genes undergoing a Polycomb (PcG)/TrxG switch during their transcriptional activation, where H3K27me3 is removed and replaced by H3K4me3 (Ringrose and Paro, 2004; Köhler and Hennig, 2010; Kuroda *et al*., 2020).

In plants, both H3K4me3 and H3K27me3 have been implicated in the control of development, but also of stress responses (Köhler and Hennig, 2010; Kleinmanns and Schubert, 2014; Engelhorn *et al*., 2017; Faivre and Schubert, 2023). Indeed, several PcG proteins are necessary for the repression of stress responses in plants growing in optimal conditions (Alexandre *et al*., 2009; Kim, Zhu and Renee Sung, 2010; Kleinmanns *et al*., 2017) while numerous TrxG members have been shown to be essential to the proper induction of stress responses (Ding, Avramova and Fromm, 2011; Song *et al*., 2021). In addition to the immediate control of stress responses, both H3K4me3 and H3K27me3 also regulate the memory of past stress episodes (Friedrich *et al*., 2018; Yamaguchi *et al*., 2021). However, the potential role of both methylation marks in the response to cold and in cold acclimation remains largely underinvestigated. Numerous *COR* genes carry H3K27me3 in the absence of cold (Vyse *et al*., 2020) and the repressive mark is lost on certain loci during cold exposure (Kwon *et al*., 2009). H3K27me3 has therefore been hypothesized to maintain the *COR* genes in a silenced state until the plant perceives low temperatures, at which point the repression is lifted through demethylation. However, previous work from our lab demonstrated that not all H3K27me3-carrying *COR* genes undergo demethylation during cold exposure (Vyse *et al*., 2020), raising questions on both the role of H3K27me3 and its removal in the control of cold responses. In order to shed more light on the putative contribution of H3K27me3 to cold acclimation, we performed a genome-wide profiling of its distribution during cold exposure. As stress-responsive genes are commonly thought to be undergoing a PcG/TrxG switch during their activation, the distribution of H3K4me3 was also examined. We uncovered a rapid redistribution of both methylation marks upon cold exposure, albeit on distinct sets of genes. By combining the epigenomic approach with a transcriptomic study, we identified a correlation between differential methylation and differential expression. However, differential methylation was not required for the transcriptional activation of *COR* genes, but might favor a higher amplitude of induction. Finally, we examined the impact of reduced H3K27me3 levels in the *clf* mutant on the cold acclimation response and could not detect any significant difference in physiological or transcriptional responses, suggesting that H3K27me3 might not participate directly in the cold response but rather in more long-term responses or to the deacclimation process. Alternatively, H3K27me3 levels may only be sufficiently reduced in *clf swn* double mutants for unmasking the role of H3K27me3 in cold acclimation.

## 2 Material and Methods

### 2.1 Plant material and growth conditions

*Arabidopsis thaliana* accession Columbia (Col0) was used as a wild type. The *clf-28* line (SALK_139371) was obtained from the Nottingham Arabidopsis Stock Centre (NASC). The primers used for genotyping are listed in Supplementary Information Table S1. The seeds were surface-sterilized, stratified in the dark at 4 °C for three days and grown on ½ MS media supplemented with Gamborg B5 vitamins (Duchefa) containing 1.5% (w/v) plant agar (Duchefa) in short day conditions (8 h light, 16 h darkness) at 20 °C for 21 days. Cold treatments were performed at 4 °C in short day conditions for 3 hours or 3 days.

### 2.2 Electrolyte leakage

Plants were grown as described previously for 21 days and placed at 4°C for three days. The freezing tolerance was then measured by electrolyte leakage assay using a protocol adapted from Hincha and Zuther (2014). Four technical replicates were performed for each biological replicate. For each sample, six temperature points were measured, using a pool of shoot tissue of five to eight seedlings. The LT50 was determined using the non-linear regression log(agonist) vs response from the GraphPad Prism version 7.0 (GraphPad Software).

### 2.3 Western Blot

100 mg of 21 day-old seedlings were harvested 4 hours after the light onset and flash-frozen in liquid nitrogen. The histones were extracted following the protocol described in Bowler *et al*. (2004) with the following modifications: the samples were resuspended in 1 mL of buffer 1. After filtration through Miracloth, the samples were centrifuged 20 min at 4000 rpm at 4 °C. The pellets were resuspended in 300 μL of buffer 2, centrifuged 10 min at 13000 rpm at 4°C and resuspended in 300 μL of buffer 3 and layered on 300 μL of clean buffer 3. After a 1 h centrifugation at 13000 rpm at 4°C, the pellets were resuspended in 100 μL of nuclei lysis buffer. The protein concentration was assessed using the Qubit protein assay (ThermoFisher Scientific) and all samples were adjusted to the same concentration using nuclear lysis buffer. The immunoblot analysis was performed as described in Hisanaga *et al*. (2023) using the following antibodies: α-H3K27me3 (C15410195 Diagenode), α-H3K4me3 (C15410003, Diagenode) and α-H3pan (C15200011 Diagenode). The imaging was performed using the Image Studio Lite software (Li-Cor, version 5.2). The intensity of the H3K27me3 and H3K4me3 signals were normalized to the intensity of the H3 signal.

### 2.4 ChIP-qPCR

1 g of 21 day-old seedlings was harvested 4 hours after the light onset. The cross-linking reaction, chromatin extraction and immunoprecipitation were performed as previously described in Vyse et al. (2020). The chromatin was incubated with 1 μg of α-H3K27me3 (C15410195 Diagenode), α-H3K4me3 (C15410003, Diagenode), α-H3pan (C15200011 Diagenode) or α-IgG (C15410206 Diagenode) antibodies. The qPCR was performed using the Takyon ROX SYBR MasterMix blue dTTP kit and the QuantStudio5 (Applied Biosystems). The primers used for the ChIP-qPCR analysis are listed in Supplementary Information Table S1.

### 2.5 ChIP-seq analysis

After DNA recovery, the DNA was purified and concentrated using the ChIP DNA Clean and Concentrator lit (Zymo Research). The libraries were prepared using the ThruPLEX DNA-seq kit (Takara Bio) and indexes from the SMARTer DNA HT Dual Index kit (Takara Bio). DNA fragments were then selected based on size using AMPure beds (Beckman Coulter). The concentration of the samples was measured using the Qubit dsDNA High Sensitivity kit and the Qubit Fluorometer (ThermoFisher Scientific) and the library quality was assessed using the High Sensitivity DNA ScreenTape and the TapeStation (Agilent). The libraries were sequenced by Novogene (UK) using a HiSeq instrument (Illumina) in 150bp paired-end mode. Two biological replicates were performed, a summary of the reads number is given in Supplementary Information Table S2.

Bioinformatic analyses were performed using Curta, the High Performance Computing of the Freie Universitaet Berlin (Bennet, Melchers and Proppe, 2020). The reads were mapped to the TAIR10 reference genome of *Arabidopsis thaliana* using Bowtie2 (Langmead and Salzberg, 2012). PCR duplicates and reads with an aligment quality MAPQ < 10 were removed using samtools rmdup and samtools view respectively (Li *et al*., 2009). The peak calling was performed using MACS2, using the broad option and a p-value threshold of 0.01 (Gaspar, 2018). Bigwig tracks were generated by pooling the two replicates and normalizing as RPKM using DeepTools bamCoverage, using a bin size of 10bp (Ramírez *et al*., 2016) and visualized using the IGV genome browser (Robinson *et al*., 2011).

Read counts for each nuclear-encoded gene (from TSS to TES) were obtained using featureCounts (Liao, Smyth and Shi, 2014) and fold changes were computed using DESeq2 (Love, Huber and Anders, 2014). A gene was considered differentially methylated if (i) it was located within a peak of the histone mark in at least one of the tested condition and (ii) it showed an absolute log2 fold change of at least 0.5. The metagenes plots were produced using deepTools (Ramírez *et al*., 2016) on the merged RPKM bigwig files, scaling all genes to 2000 bp and examining a region starting 500 bp upstream from the TSS and ending 500 bp downstream form the TES.

### 2.6 RT-qPCR

100 mg of seedlings were harvested 4 hours after light onset and flash frozen in liquid nitrogen. After grinding to a fine powder, total RNA was extracted using the innuPREP Plant RNA kit (Analytik Jena). Samples were treated with DNaseI (ThermoFisher Scientific) and cDNA was synthesized using the RevertAid Reverse Transcriptase kit (ThermoFisher Scientific). The qPCR was performed using the Takyon ROX SYBR MasterMix blue dTTP kit and the QuantStudio5 (Applied Biosystems). The primers used for the RT-qPCR analysis are listed in Supplementary Information Table S1. The Ct values were normalized by subtracting the mean of three housekeeping genes (*ACTIN2*, *PDF* and *TIP41*) from the Ct value of each gene of interest (ΔCt). Transcript abundance was expressed as 2^−ΔCt^.

### 2.7 RNA-seq analysis

RNA samples were extracted and DNaseI-treated as previously described. The libraries were prepared using poly-A enrichment by Novogene (UK) and the sequencing was performed on the NovaSeq 600 platform (Illumina) in 150bp paired-end mode. Three biological replicates were analysed and a summary of the reads number is given in Supplementary Information Table S2.

Bioinformatic analyses were performed using Curta, the High Performance Computing of the Freie Universitaet Berlin (Bennet, Melchers and Proppe, 2020). The reads were mapped to the reference genome of Arabidopsis thaliana (TAIR10) using STAR (Dobin *et al*., 2013), using a minimum and maximum intron size of 60 and 6000 bases respectively. The counting was performed using featureCounts (Liao, Smyth and Shi, 2014), using only reads with an alignment score superior to 10. The differential expression analysis was performed using the DESeq2 package (Love, Huber and Anders, 2014). A gene was considered to be differentially expressed (DEG) if it presented an absolute log2 fold change of at least 1 and a Benjamini-Hochberg adjusted p-value inferior to 0.05. As the differences in expression were correlated to the differences in histone methylation levels, only nuclear-encoded DEGs were retained in the analysis.

### 2.8 Statistics and data visualization

Unless stated otherwise, statistical analyses and plots were generated using R or GraphPad Prism (GraphPad Software). Normal distribution was tested using the Shapiro-Wilks’ method. For normally distributed data, ANOVA tests and any post-hoc tests were performed using the agricolae package (de Mendiburu and Yaseen, 2020).

Gene ontology enrichment analyses were performed in RStudio using the topGO package (Alexa and Rahnenfuhrer, 2021), the TAIR10 annotation and the gene-GO term relationships from the org.At.tair.db package, version 3.17.0 (Carlson, 2019).

## 3 Results

### 3.1 H3K4me3 and H3K27me3 undergo differential methylation upon short cold exposure

To determine whether cold exposure triggers genome-wide changes in the levels of H3K27me3 and H3K4me3, a Western-Blot was conducted on plants exposed to 4°C for three hours or three days (Figure 1A and B). For both chromatin marks, no genome wide changes could be detected at the time points tested here. However, previous studies indicated that H3K27me3 is removed from certain loci upon cold exposure while H3K4me3 was shown to be accumulated at others, suggesting that both marks might undergo differential methylation in a loci-specific manner that does not lead to changes detectable at the genome wide scale (Kwon *et al*., 2009; Miura, Renhu and Suzaki, 2020; Vyse *et al*., 2020). To assess this possibility, an epigenome profiling of the distribution of H3K4me3 and H3K27me3 was performed at the same time points described above. In total, 13829, 14152 and 14430 H3K4me3 peaks were detected in naïve, 3h and 3d samples respectively while 5753, 5665 and 5802 H3K27me3 peaks were detected in those same samples. These peaks largely overlapped for the individual marks, indicating that short cold exposure did not lead to a substantial redistribution of the chromatin methylation marks investigated here. In order to detect lower magnitudes of methylation levels changes, reads mapped between the transcription start site (TSS) and transcription end site (TES) of genes targeted by each methylation mark were counted and normalized for each condition (Figure 1C to F). The correlation plots indicated that H3K4me3 is accumulated after three hours of cold exposure while after three days, this tendency mostly disappeared (Figure 1C and E). On the other hand, H3K27me3 correlation plots displayed an accumulation of the mark at both time points (Figure 1D and F). The differentially methylated genes were identified as genes targeted by the respective mark (i.e. covered by a peak in a least one condition) and showing an absolute log2FC of the normalized counts of at least 0.5. The complete list of differentially methylated (DM) genes can be found in Supplementary Table 3. Consistent with the general trend observed on the correlation plots, more genes were found to significantly gain H3K4me3 or H3K27me3 than losing it. 3619 and 2309 DM genes were identified for H3K4me3 after three hours and three days of cold treatment, respectively, while H3K27me3 differential methylation was detected only on 735 and 922 genes, respectively. This substantial disparity in the number of DM genes between H3K4me3 and H3K27me3 can be largely explained by the fact that H3K4me3 targets a broader proportion of genes than H3K27me3 (17366 vs 8128): between 13 and 20% of H3K4me3 targets are differentially methylated while only 9 to 11% of H3K37me3 targets undergo changes during cold exposure.

**Figure 1:**
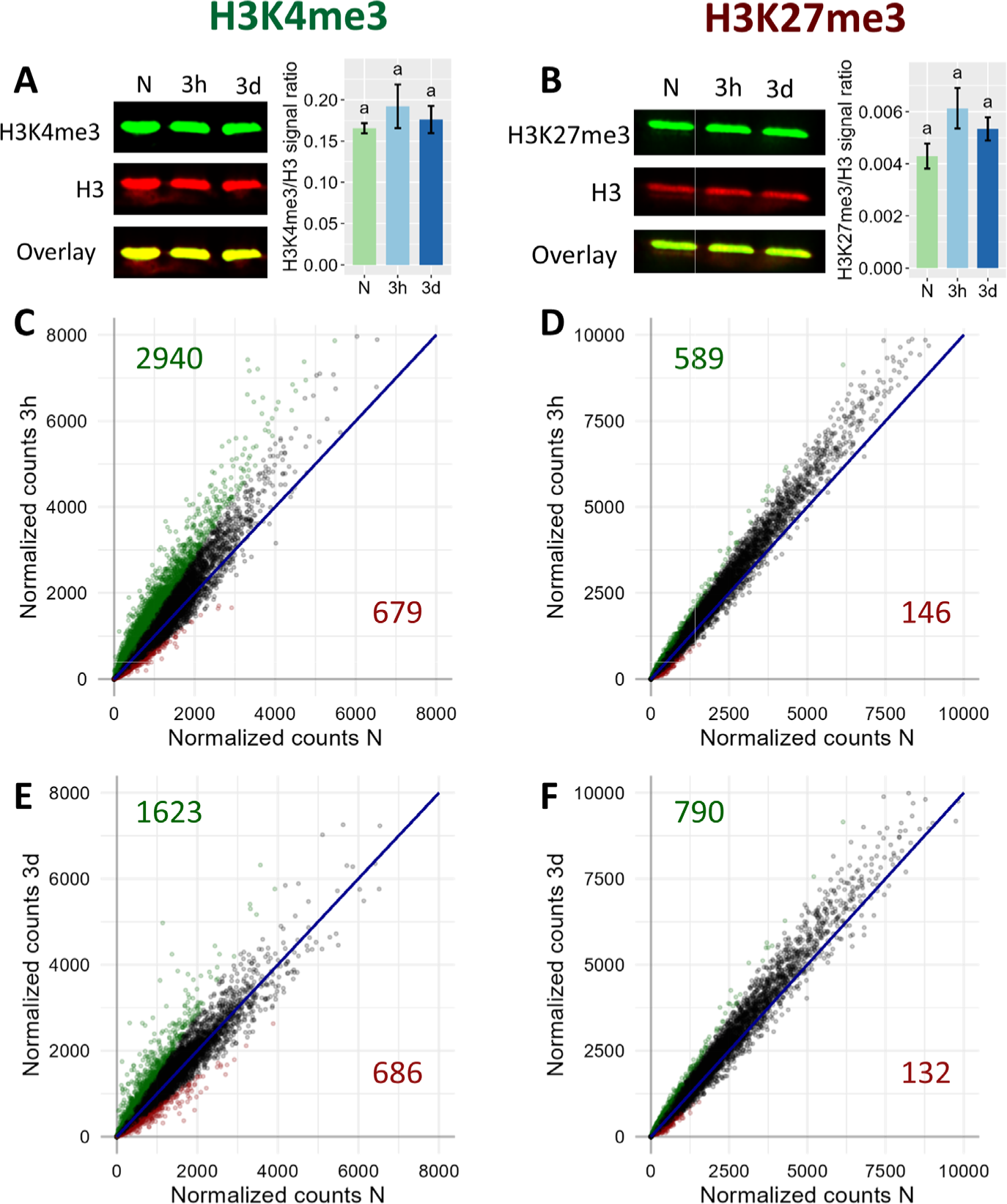
Genome-wide dynamics of H3K4me3 (left) and H3K27me3 (right) upon cold exposure. Plants were grown for 21 days at 20°C (N) and then exposed to 4°C for three hours (3h) or three days (3d). **(A)** and **(B)** Global levels of H3K4me3 and H3K27me3, respectively, as measured by Western Blot. The membrane images show the signal of the histone methylation mark in green, of total histone 3 in red and the overlay of both signals in yellow. The bar charts on the right of the membrane images display the modification/H3 signal ratio of four independent biological replicates. Significance was tested by one-way ANOVA followed by a Tukey post-hoc test (α = 0.05). Identical letters indicate no significant difference. **(C)** to **(F)** Correlation plot of H3K4me3 (**(C)** and **(E)**) and H3K27me3 (**(D)** and **(F)**) levels on genes targeted by the respective mark after 3h (**(C)** and **(D)**) or 3d (**(E)** and **(F)**) of cold exposure. Each point represents a gene targeted by the respective mark. Reads were counted over the gene body and were normalized to library size using DESeq2 (See Material and Methods). Genes showing a log2 fold change of the respective mark smaller than -0.5 are displayed in red, while genes showing log2 fold change of at least 0.5 are displayed in green. Total number of genes satisfying these criteria are indicated in red in the lower right quadrant and green in the upper left quadrant respectively.

While the proportion of DM genes is not strikingly different between H3K4me3 and H3K27me3, the magnitude of the changes differs significantly, with H3K4me3 DM genes presenting higher absolute fold change values than H3K27me3 DM genes (Figure 1C to F, Supplementary Figure 1). These observations were confirmed when examining the levels of both methylation marks at specific loci (Figure 2A): the changes of H3K4me3 were drastic, leading to peaks appearing (*CBF3*, *LTI30* and *COR15A*) or disappearing (*HSP90.1*). The changes were prominently located just downstream of the TSS, consistent with the known localization of H3K4me3, whose peaks usually center around the TSS of its target genes, and were more pronounced after three days than after three hours (Supplementary Figure 1A). On the other hand, while H3K27me3 loss led to the almost-complete loss of peaks at certain loci such as *LTI30*, it was more limited on other such as *COR15A*, where the H3K27me3 peaks were still visible after three days of cold treatment. Genes gaining H3K27me3 showed moderately increased levels of the repressive mark on the sides of the original peak (*END1* and *AT5G43570*). The variations in H3K27me3 occurred on the whole gene body of the DM genes and were more pronounced in the case of loss than of gain (Supplementary Figure 1B). Overall, even short cold exposure times of three hours were sufficient to trigger significant alteration of the level of both methylation marks on thousands of loci.

**Figure 2:**
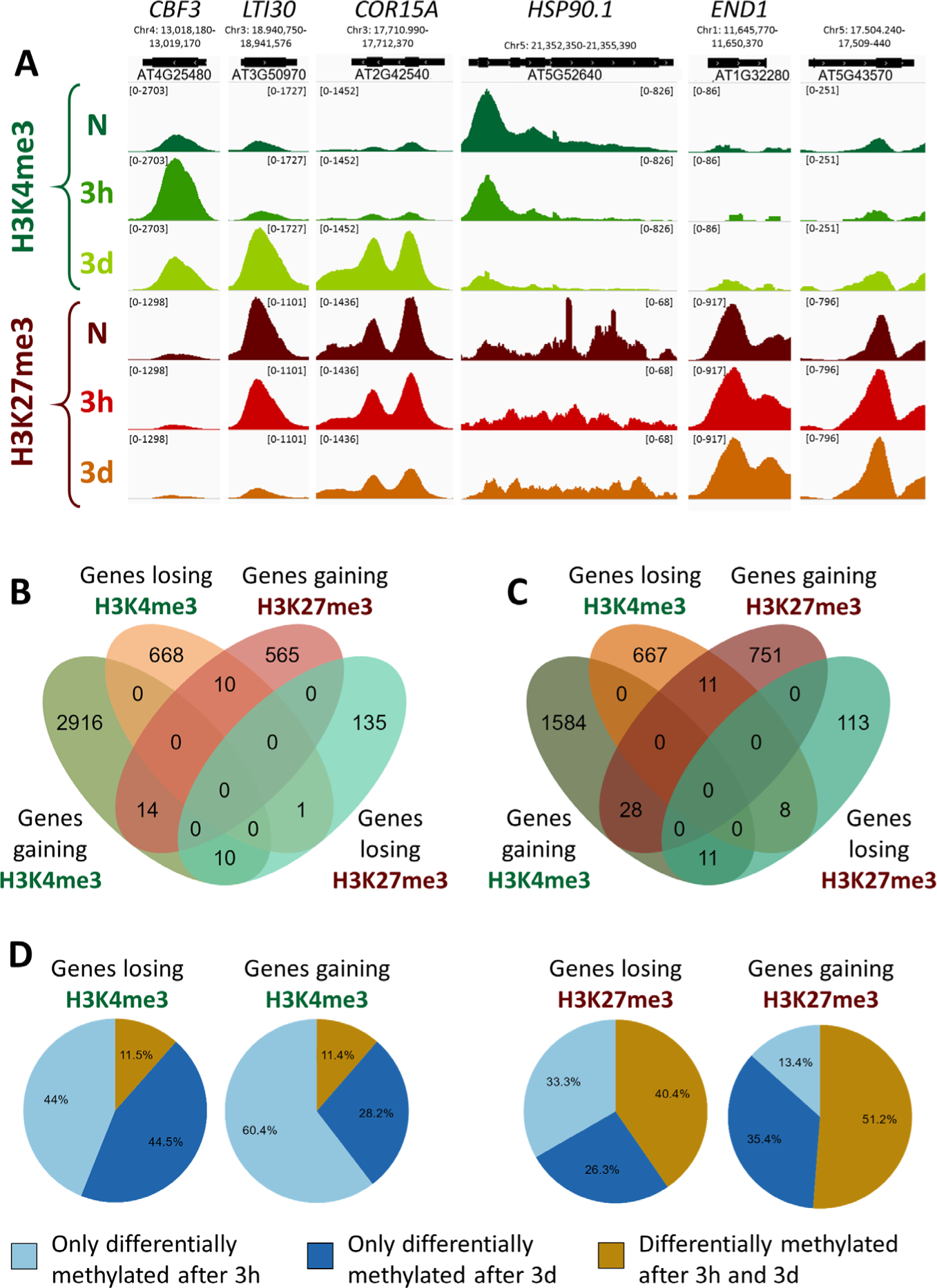
Characterization of differentially methylated genes. A differentially methylated gene is defined as a gene targeted by H3K4me3 or H3K27me3, respectively, and showing an absolute log2 fold change of the respective methylation level of at least 0.5. **(A)** Genome browser views of H3K4me3 and H3K27me3 ChIP-seq signals at selected differentially methylated genes, in naïve plants (N) or plants exposed to 4°C for 3h or 3d. The numbers in bracket at the top of each track indicate the scale of that track in reads per million per bin. **(B)** and **(C)** Venn diagrams showing the overlaps of differentially methylated genes for both histone methylation marks after 3h or 3d in the cold respectively. **(D)** Pie charts indicating the percentage of differentially methylated genes being specifically regulated at a single time point or at both time points examined, for each histone mark and direction of differential methylation.

### 3.2 H3K4me3 and H3K27me3 differential methylation occurs on stress responsive and developmental genes, respectively

Overlapping the sets of DM genes at each time point revealed that only a minor proportion of them undergo a change in both H3K4me3 and H3K27me3 levels, totaling 35 genes at the 3h time point and 58 at the 3d time point (Figure 2B and C). As those marks are commonly described as antagonists, genes differentially methylated for both marks would be expected to display opposite changes. However, there are only slightly fewer genes showing same direction changes than opposite (10 vs 25 at 3h, 22 vs 36 at 3d), suggesting that the loss of one mark does not entail a gain of the other and vice versa. As the DM gene sets of H3K4me3 and H3K27me3 displayed such a reduced overlap, we hypothesized that H3K4me3 and H3K27me3 differential methylation might serve distinct purposes. To explore this hypothesis, a gene ontology (GO) term analysis for biological function was performed on each DM gene set (Supplementary Figures 2 and 3, Supplementary Table 5). Genes gaining H3K4me3 during a cold treatment were enriched for terms related to the cold response, cold acclimation and freezing tolerance as well as terms linked to the response to other abiotic and biotic stresses (water deprivation, hypoxia, fungus) (Supplementary Figure 2A and C). After three hours of cold exposure, genes losing H3K4me3 were enriched for terms related to protein refolding and chromatid cohesion, while after three days the set showed an enrichment for development and photosynthesis related terms (Supplementary Figure 2B and D). Few terms were found to be enriched among the genes losing H3K27me3, which might be due to the smaller size of the sets (Supplementary Figure 2B and C). Some terms related to stress responses were identified (response to salicylic acid and to fungus) but surprisingly, no term associated to the cold response was found to be enriched. Genes gaining H3K27me3 upon cold exposure were mostly enriched for development related terms (Supplementary Figure 3A and C). H3K4me3 and H3K27me3 differential methylation therefore occur on different sets of genes, with H3K4me3 DM mostly targeting stress responsive genes and H3K27me3 DM developmental genes. This could suggest that differential histone methylation holds a distinct role in the cold response depending on the specific mark.

To determine whether the methylation changes triggered by cold exposure were stable over time or dynamic, their persistence was examined by computing the percentage of genes differentially methylated at both time points (Figure 2D). In the case of H3K4me3, only 11% of the DM genes were identified at both time points, indicating that the variations in the level of the active mark were rather transient. On the contrary, 40 to 50% of H3K27me3 DM genes displayed a change at both time points, revealing H3K27me3 changes to be more stable over time than those of H3K4me3. Taken together with the results of the GO analysis and the small overlap between the genes which are DM for H3K4me3 and H3K27me3, this suggests that H3K4me3 and H3K27me3 differential methylation might serve distinct purposes.

### 3.3 Differential methylation partially correlates with differential expression

As H3K4me3 and H3K27me3 are commonly described as favoring and silencing transcription, respectively, we hypothesized that the changes in the levels of those two chromatin marks might associate with differences in the transcriptional activity of the underlying genes. A transcriptome analysis was therefore performed on the same seedlings used for the epigenome investigations, leading to the identification of the nuclear-encoded genes up- and down-regulated after three hours or three days of cold exposure (Supplementary Table 4). After three hours of cold treatment, no correlation between the changes in H3K4me3 levels and the changes in expression could be detected (Figure 3A), while a weak negative correlation was observed for H3K27me3 (Figure 3B). However, after three days, the changes in expression were positively and negatively correlated with the variations in H3K4me3 and H3K27me3, respectively (Figure 3C and D), indicating that genes up-regulated by a cold treatment were more likely to gain H3K4me3 and/or lose H3K27me3. Since those correlations were seen after three days of cold exposure but not (or to a lesser extend) after three hours, it is likely that the two phenomenon (differential expression and differential methylation) occur at a different pace. Indeed, after three days at 4°C, the plants are accustomed to the cold and cold acclimation can already be detected at a physiological level, while after only three hours, the plant is only starting its acclimation process and not all responses are fully accomplished yet (Calixto *et al*., 2018; Zuther *et al*., 2019). To try and decipher whether chromatin or expression changes first, the correlations analyses were repeated across time points (Figure 3E). The early methylation changes did not strongly correlate with the late expression changes. However, early expression changes correlated positively and negatively with the variations in H3K4me3 and H3K27me3 levels, respectively. Those results collectively suggest that the transcriptional activity of a gene is modulated ahead of its chromatin methylation status, but more precise time-course experiments would be required to fully confirm this observation.

**Figure 3:**
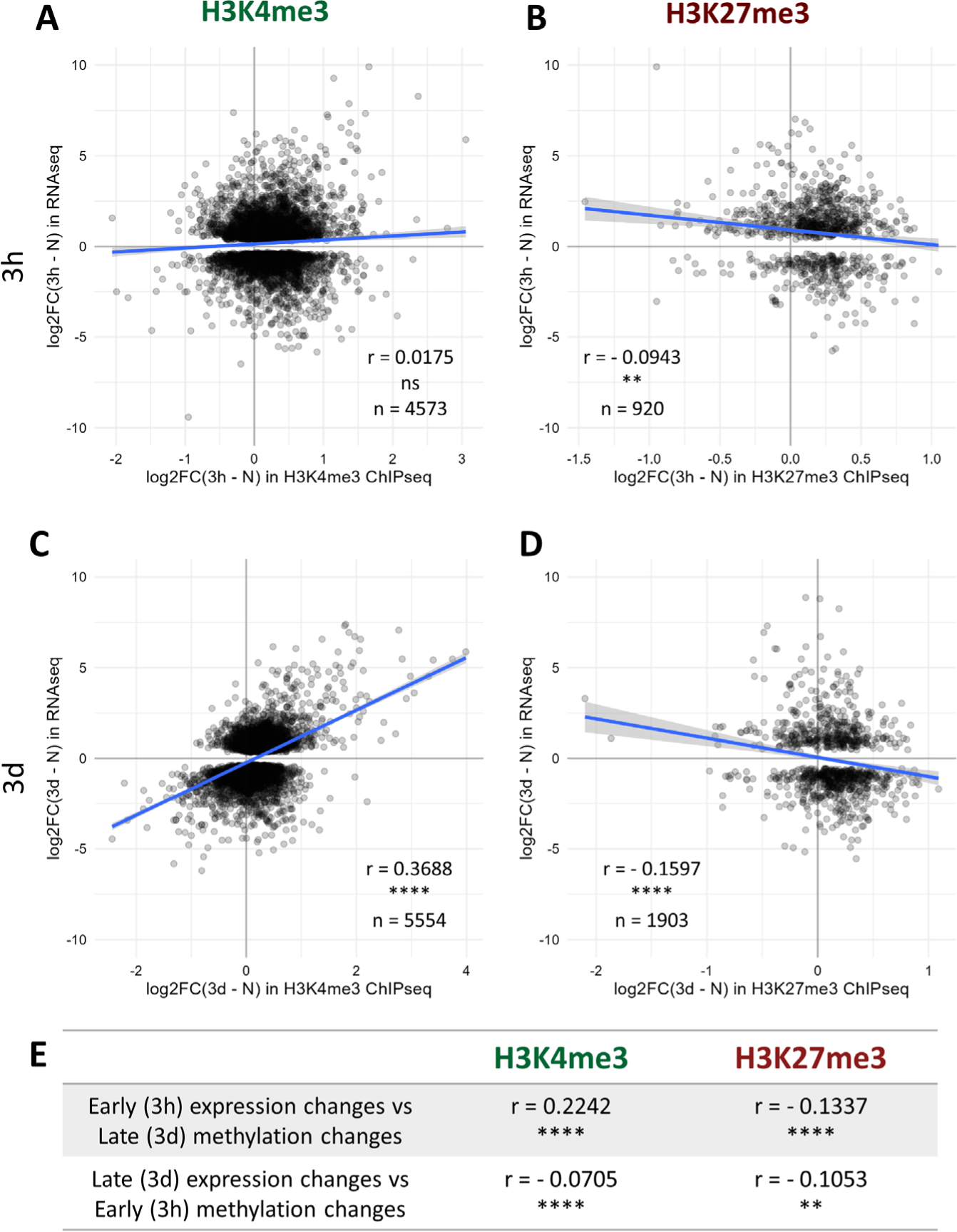
Correlation between histone methylation and expression changes upon cold exposure. Plants were grown for 21 days at 20°C (N) and then exposed to 4°C for three hours (3h) or three days (3d). H3K4me3 and H3K27me3 levels were measured by ChIP-seq while the changes in expression were detected by an RNA-seq conducted on RNA isolated from the same seedlings. Correlation between changes in expression and changes in H3K4me3 (**(A)** and **(C)**) or H3K27me3 (**(B)** and **(D)**) levels after 3h (**(A)** and **(B)**) or 3d (**(C)** and **(D)**) of cold exposure. For each graph, the X axis denotes the log2 fold change in methylation signal over the whole gene body at the respective time point compared to non-cold treated plants while the Y axis shows the log2 fold change in expression for the same comparison. Only genes which are differentially expressed at the considered time point, i.e. present an absolute log2 fold change >= 1 and a p-adj < 0.05, and are targeted by the respective mark are shown on the scatterplot, their number is indicated as n. The correlation analyses were performed using the Spearman method, the correlation coefficient is indicated as r. ns indicates a non-significant correlation, ** denotes a p-value < 0.01 and **** a p-value < 0.0001. **(E)** Table summarizing the correlation between changes in expression and in methylation levels across time points, performed as described above.

Although significant correlations between methylation and expression changes could be detected, their magnitude was relatively modest. The lists of significantly differentially expressed (DE) and DM genes exhibited a moderate overlap (Supplementary Figure 6), indicating that differential methylation is not required for differential expression and that differential expression does not necessarily result in differential methylation. Whether differential methylation contributes, even partially, to the transcriptome reprogramming remains unelucidated. In order to examine whether it might facilitate the induction of cold responsive genes, the transcriptional activity of non DM and DM genes was compared for each chromatin mark (Figures 4 and 5). Out of the 17366 genes detected as carrying H3K4me3 at any time point of the stress regimen, 1714 were up-regulated upon cold treatment (Figure 4A). The majority (57%) of these genes did not undergo differential methylation, as observed on the Venn diagrams (Supplementary Figure 6), while the levels of H3K4me3 increased on 37% of the genes and decreased for 6% of them, consistent with the correlation analyses (Figure 3). The genes undergoing differential methylation had slightly lower H3K4me3 levels than non-DM genes prior to cold exposure (Figure 4D). This was associated with a lower basal expression of genes losing H3K4me3, but no difference in expression in naïve conditions could be seen between non DM genes and genes gaining H3K4me3 upon cold exposure (Figure 4B). During a cold stress, the expression of genes gaining H3K4me3 increased significantly more than those of non DM and genes losing H3K4me3 and reached higher overall expression levels (Figure 4B and C). There was however no difference in the fold change of gene expression between non DM and genes losing H3K4me3. This suggests that H3K4me3 gain, while not strictly necessary for gene activation, might facilitate it, leading to a higher magnitude of induction.

**Figure 4:**
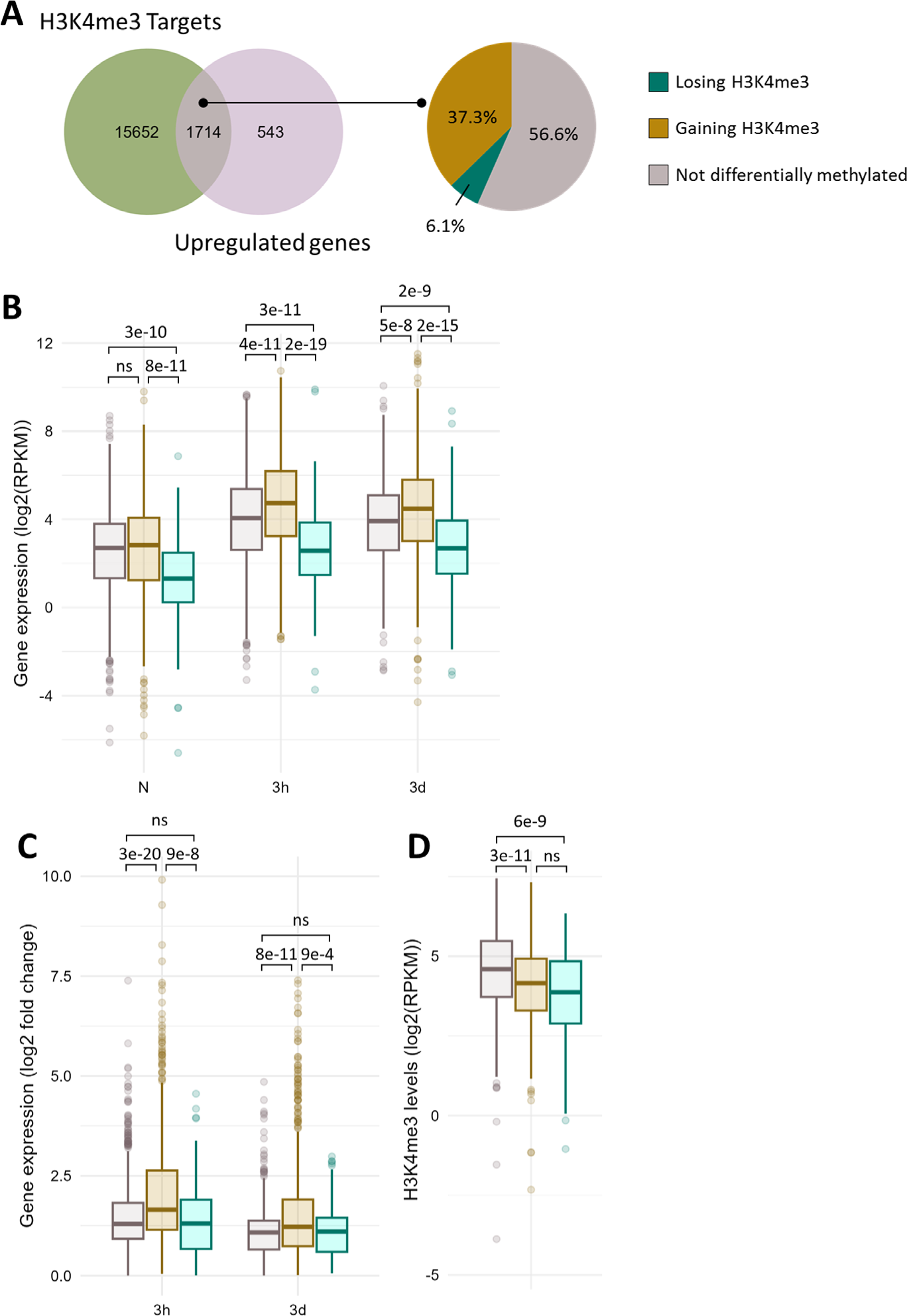
A subset of cold induced genes gain H3K4me3 upon cold exposure. **(A)** Venn diagram showing the overlap between the genes carrying H3K4me3 and the genes induced at any time point during cold exposure (left panel). Pie chart showing the percentage of genes gaining or losing H3K4me3 at any time point during cold exposure among the 1714 genes which are induced by cold and carry H3K4me3 (right panel). **(B)** Box plot showing the distribution of gene expression during cold exposure for the three gene categories listed in (A). Gene expression is shown as log2 of the RPKM (Read Per Kilobase per Million mapped read). **(C)** Box plot showing the distribution of log2 fold change in gene expression after 3h and 3 days of cold exposure for the three gene categories listed in (A). **(D)** Box plot showing the distribution of H3K4me3 levels as RPKM for the three gene categories listed in (A). The p-value were computed using a two-sided Wilcoxon rank-sum test.

**Figure 5:**
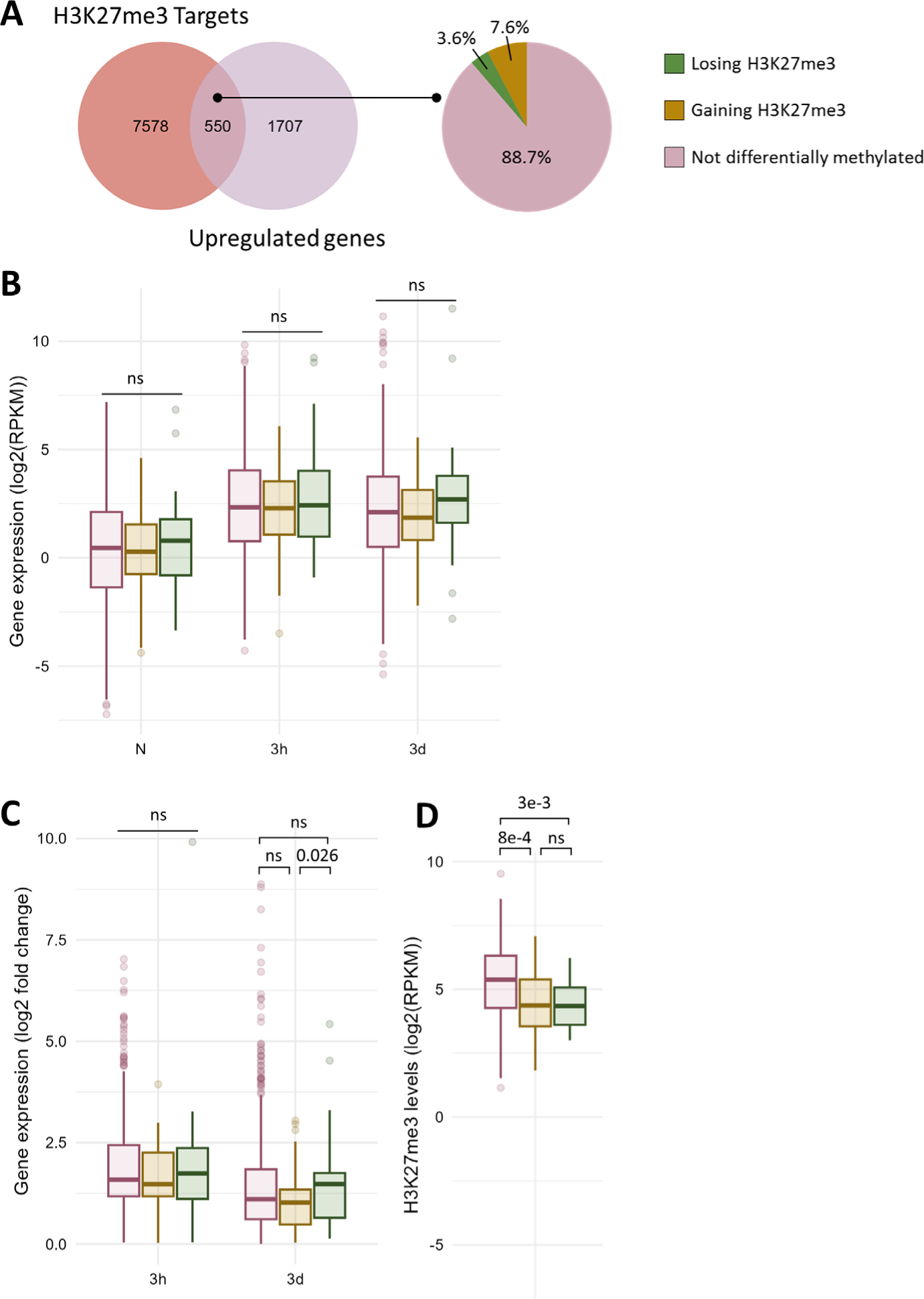
Only a fraction of cold induced genes carrying H3K27me3 undergo differential methylation. **(A)** Venn diagram showing the overlap between the genes carrying H3K27me3 and the genes induced at any time point during cold exposure (left panel). Pie chart showing the percentage of genes gaining or losing H3K27me3 at any time point during cold exposure among the 550 genes which are induced by cold and carry H3K27me3 (right panel). **(B)** Box plot showing the distribution of gene expression during cold exposure for the three gene categories listed in (A). Gene expression is shown as log2 of the RPKM (Read Per Kilobase per Million mapped read). **(C)** Box plot showing the distribution of log2 fold change in gene expression after 3h and 3 days of cold exposure for the three gene categories listed in (A). **(D)** Box plot showing the distribution of H3K27me3 levels as RPKM for the three gene categories listed in (A). The p-value were computed using a two-sided Wilcoxon rank-sum test.

8128 genes have been detected as carrying H3K27me3 in at least one time point during the stress regiment, of which 550 were induced by cold (Figure 5A). Only 3.6% of those genes lost H3K27me3 during cold exposure, confirming that H3K27me3 is not an obstacle to gene induction (Vyse *et al*., 2020). Surprisingly, a higher proportion (7.6%) showed an increase in H3K27me3 levels. Both genes gaining or losing H3K27me3 upon cold treatment had lower H3K27me3 levels in naïve conditions compared to non DM H3K27me3 targets (Figure 5D). However, this difference in the levels of the repressive mark was not associated with a difference in expression in naïve conditions (Figure 5B). The expression of non DM and DM genes remained similar upon cold exposure, but the genes losing H3K27me3 showed a higher fold change of expression after three days of cold treatment compared to genes which gained H3K27me3 (Figure 5B and C). This suggests that H3K27me3 loss might also contribute to the amplitude of induction.

### 3.4 Reduced levels of H3K27me3 do not impact the cold stress response

While some H3K27me3 targets which are induced by cold showed a reduction in the level of this mark during a cold treatment (such as *LTI30* and *COR15A*), others did not show any differential methylation (*GOLS3*, *WRKY40*). To examine whether H3K27me3 might hold different roles on those two types of genes, their transcriptional activity during a cold treatment was monitored in the H3K27 methyltransferase mutant *curly leaf* (*clf*) (Figure 6). In *clf*, H3K27me3 levels were reduced by around 50% on those cold-responsive genes, suggesting that, while they are targeted by CLF, a second methyltransferase (likely SWN) is also able to deposit H3K27me3 at those loci (Figure 6A).

**Figure 6:**
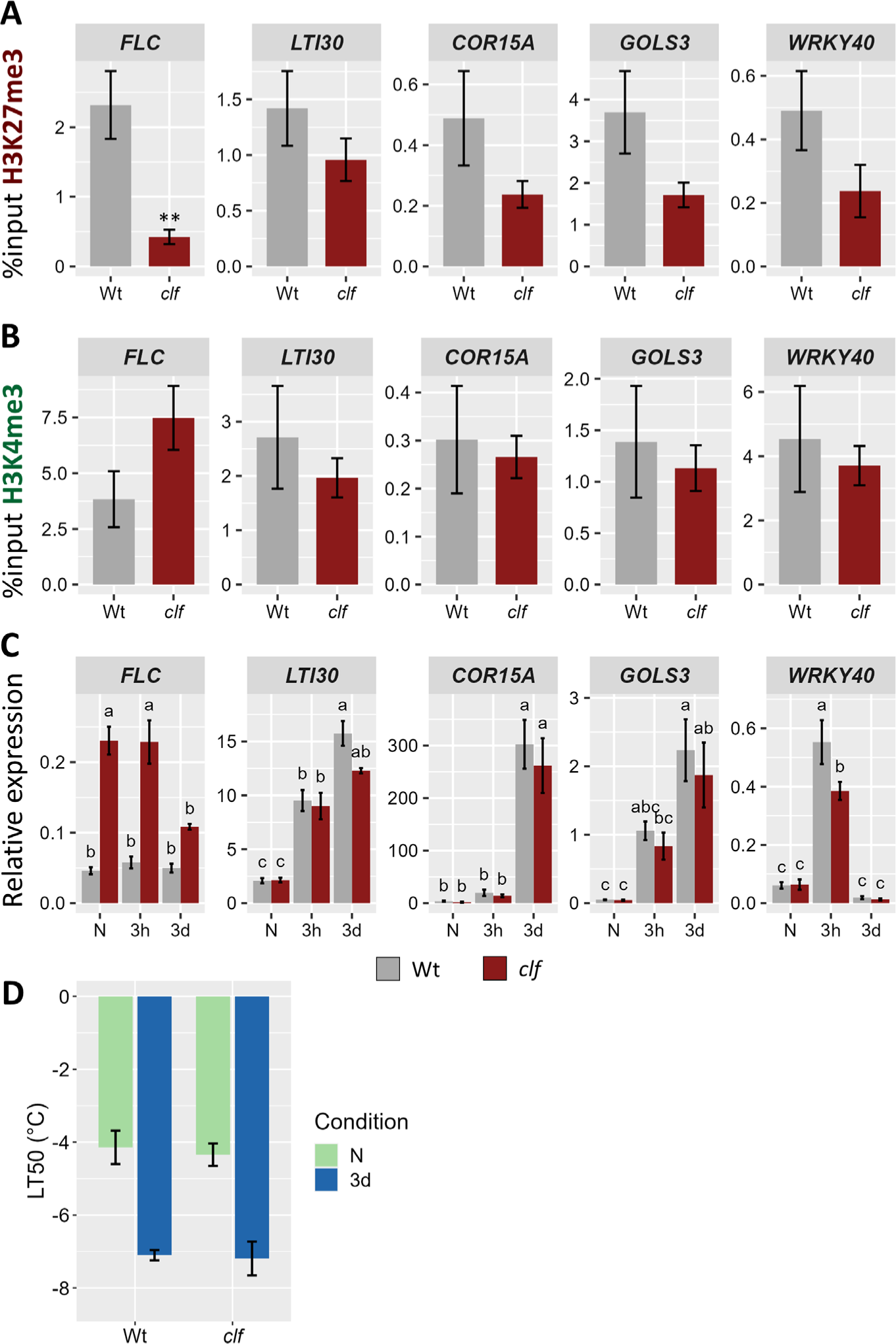
Reduced levels of H3K27me3 do not impact the transcriptional activity of cold-induced genes. **(A)** and **(B)** H3K27me3 **(A)** and H3K4me3 **(B)** levels on five H3K27me3-targeted genes in 21 day-old Wt and clf mutant plants grown at 20°C. After cross-linking, chromatin was extracted and precipitated using H3K27me3 and H3K4me3 antibodies, respectively. The purified DNA was amplified by quantitative PCR. Results are presented as %input. Error bars indicate the sem of four biological replicates. Significance was tested using t-test, ** indicates a p-value < 0.01. **(C)** Relative expression level of five H3K27me3-targeted genes in 21 day-old Wt and clf mutant plants grown at 20°C (N) and exposed to 4°C for three hours (3h) or three days (3d). Transcript levels were measured by RT-qPCR and normalized to three internal controls (TIP41, ACTIN2 and PDF). Error bars indicate the sem of three biological replicates. Significance was tested by two-way ANOVA followed by a Tukey post-hoc test (α = 0.05). Identical letters indicate no significant difference. All primer sequences used for this experiment can be found in Supplemental Table 1. **(D)** Freezing tolerance of clf mutant before or after cold acclimation, measured by electrolyte leakage assay. Plants were grown for 21 days at 20°C (N) and then exposed to 4°C for three days (3d). Error bars represent the sem of three biological replicates. Statistical significance was assessed by 2-way ANOVA followed by a Dunnett post hoc test, no significant difference was found.

Interestingly, the levels of H3K27me3 in the *clf* mutant in naïve conditions is similar to that observed after three days of cold exposure in wild-type plants (data not shown). Despite the reduced H3K27me3 levels, the expression of those genes was not altered in *clf*, neither in naïve conditions nor after a cold treatment (Figure 6C). Reduced levels of H3K27me3 did not impact the basal level of expression nor the speed or magnitude of induction. On the other hand, *FLC*, whose H3K27me3 were significantly reduced in *clf*, displayed a higher expression in this mutant in all the examined conditions. This increased expression was associated with elevated H3K4me3 levels in *clf* while the levels of this mark remained constant on the other genes (Figure 6B). Both the basal and the acquired freezing tolerance of the *clf* mutant were measured during an electrolyte leakage assay (Figure 6D). No significant difference to wild-type could be observed, confirming that reduced H3K27me3 levels do not impact cold tolerance. Altogether, these data reject the simplistic model whereby a reduction of H3K27me3 would directly lead to increased H3K4me3 levels and to the transcriptional activation of previously silenced genes.

## 4 Discussion

### 4.1 Chilling stress alters the levels of both H3K4me3 and H3K27me3 at specific loci

Low temperatures are known to alter the distribution and levels of both H3K4me3 and H3K27me3 in the genome of *Arabidopsis thaliana* (Xi *et al*., 2020). However, studies have so far either focused on long cold treatments, with the aim of investigating vernalization, or examined only a handful of loci (Kwon *et al*., 2009; Vyse *et al*., 2020; Xi *et al*., 2020). The potential contribution of histone methylation to the response to cold stress therefore remains unelucidated. Here, we attempt to shed some light on this question by performing a genome-wide investigation of H3K4me3 and H3K27me3 dynamics after short (three hours or three days) 4°C treatments. While H3K27me3 was shown to be accumulated in Arabidopsis growing in moderate heat (Kim *et al*., 2023), Western Blots did not reveal drastic changes in the levels of either mark upon cold exposure (Figure 1A and B), similarly to what was observed during chilling stress in grapevine leaves (Zhu *et al*., 2023). In order to gain a more detailed view on potential changes, ChIP-seq were performed. The distributions of both H3K4me3 and H3K27me3 were not dramatically altered upon cold treatment: the number of peaks and of genes carrying the marks were sensibly the same in all conditions. This is in stark contrast to the consequences of cold treatment in *Oryza sativa*, where only 38% of genes enriched in H3K27me3 were common to the naïve and cold-stress conditions (Dasgupta *et al*., 2022). However, it led to many local changes in the levels of both H3K4me3 and H3K27me3, with around 5 300 and 1 100 genes showing an absolute log2 fold change of at least 0.5, respectively (Figure 1C to F). Differentially methylated genes were already detected after only three hours of cold treatment, indicating that this process happens on a time scale similar to that of differential expression (Calixto *et al*., 2018). For both marks, differential methylation was skewed towards a gain of the modification, while after 40 days of cold treatment, Xi et al. (2020) observed a trend of gain for H3K27me3 and of loss for H3K4me3, hinting that varying lengths of cold treatment might impact the distribution of histone marks differently. Significantly more genes underwent H3K4me3 differential methylation than H3K27me3, but once reported to the total number of genes targeted by each mark, the difference was not substantial anymore. However, H3K4me3 changes were, on average, of a larger magnitude than those observed for H3K27me3 (Figure 1C to F, Supplementary Figure 1).

### 4.2 The induction of cold stress responsive genes does not rely on a PcG-TrxG switch

Both H3K4me3 and H3K27me3 differential methylation correlated with differential expression, especially after a longer (three days) cold exposure (Figure 3). Genes induced by cold generally displayed a gain of H3K4me3 and/or a loss of H3K27me3. However, very little overlap between H3K4me3 and H3K27me3 DM genes was observed, refuting the simplistic model of stress-responsive genes transitioning from a silenced H3K27me3 chromatin to an active form enriched in H3K4me3 during their transcriptional activation. This was further confirmed by examining the levels of H3K4me3 on cold-inducible genes in the *clf* mutant (Figure 6): while their H3K27me3 status was reduced, no significant difference in H3K4me3 could be observed, indicating that H3K27me3 is not automatically replaced by H3K4me3. Such a PcG/TrxG switch has been demonstrated for transcriptional activation during development (Engelhorn *et al*., 2017), but in this context, the expression of the gene is altered indefinitely. By contrast, stress responses only require a transient adjustement of the transcriptional activity. It is therefore possible that the chromatin status of cold responsive genes is not as dramatically altered, to allow for reversion to the initial state once the stress subsides. Instead of a H3K27me3-to-H3K4me3 switch, H3K4me3 and H3K27me3 differential methylations appear to be mostly independent from one another, suggesting that they might hold very distinct functions. Indeed, the GO analyses uncovered that distinct categories of terms were enriched for H3K4me3 and H3K27me3 DM genes (Supplementary Figures 2 and 3). Furthermore, the correlation between differential methylation and differential expression was stronger for H3K4me3 than H3K27me3. This is consistent with a previous study from Engelhorn *et al*. (2017) on the floral transition, which reported H3K4me3 to be a stronger predictor of transcriptional changes than H3K27me3. The levels of the active mark were also altered prior to the ones of its silencing counterpart during seasonal oscillations (Nishio *et al*., 2020). In the present study, while both marks already displayed variations after only three hours of cold exposure, only about 11% of H3K4me3 changes were detected both after three hours and three days of cold treatment, suggesting that they are mostly transient (Figure 2D). On the contrary, the majority of H3K27me3 changes were shared by both time points, indicating a higher stability of H3K27me3 modifications. This is consistent with previous analyses of H3K4me3 and H3K27me3 dynamics in HeLa cells, which reported H3K4me3 as having a faster turn-over and re-establishment speed than H3K27me3 (Zheng *et al*., 2014; Alabert *et al*., 2015; Reverón-Gómez *et al*., 2018). Mathematical modelling demonstrated that chromatin marks with slower dynamics are more robust against rapidly fluctuating environmental conditions, as the signal has to persist longer for a new equilibrium for the level of the mark to be reached (Berry, Dean and Howard, 2017). The different dynamics of H3K4me3 and H3K27me3 could therefore confer them different responsiveness to environmental variations, with H3K4me3 contributing to the immediate stress response to lower temperature (as suggested by the enrichment of abiotic and biotic stress-response related GO terms) and H3K27me3 mediating more long term responses such as developmental adaptations (Supplementary Figures 2 and 3).

### 4.3 Role of differential methylation in gene regulation

It has been reported that cold-inducible H3K27me3 targets lose the repressive mark upon induction (Kwon *et al*., 2009), but in a previous work, we demonstrated that loss of H3K27me3 is not required for induction (Vyse *et al*., 2020). This new genome wide analysis confirms our prior report and refutes the idea that H3K27me3 is an absolute obstacle for transcriptional activation of cold-responsive genes. To further dissect the potential role of cold-induced H3K27me3 loss on those genes, we used the *clf* mutant, in which many cold-responsive genes present a reduced H3K27me3 status. The reduced H3K27me3 levels in the *clf* mutant did not lead to a change in the basal expression of the genes investigated here (Figure 6C), suggesting that additional factors are required for their transcriptional activation, likely transcription factors such as the CBFs. Similar observations were made by Liu *et al*. (2014), where the absence of a functional CLF and therefore the reduction of H3K27me3 at drought inducible genes did not trigger their induction in naïve conditions. However, the authors observed a higher magnitude of induction upon stress exposure, which was not detected in the present study. It is therefore likely that H3K27me3 holds a different function in the response to drought and in the response to cold. Instead, the silencing mark might control the induction speed of the genes: reduced H3K27me3 status has been reported as allowing a faster transcriptional activation in the case of camalexin biosynthesis genes during pathogen infection (Zhao *et al*., 2021). This does not seem to hold true for cold-inducible genes: the expression levels in Wt and *clf* were comparable both after three hours and three days of cold exposure, suggesting that lower H3K27me3 status does not lead to a faster induction. However, more detailed time-course transcriptomic experiments would be required in order to reach a definite conclusion. Interestingly, the levels of H3K27me3 on cold inducible genes in the *clf* mutants are similar to those observed after three days of cold exposure. The lack of higher or faster induction of those genes upon cold stress in *clf* is therefore consistent with observations from Kwon et al. (2009), where a persisting cold-induced lower H3K27me3 status did not lead to an altered expression of the genes upon cold re-exposure. Furthermore, H3K27me3 does not appear to directly contribute to the regulation of the cold stress response (at least for the tested conditions), as *clf* mutants also did not show an altered basal or acquired freezing tolerance compared to wild-type (Fig 6D). Instead, H3K27me3 might contribute to the regulation of deacclimation or memory processes, only affecting transcriptional activity after the cold episode subsides. In addition, many development-related terms were identified in the gene sets gaining H3K27me3, suggesting that they might be down-regulated upon cold exposure. However, when performing a GO term analysis on the lists of genes differentially expressed after three hours and three days of cold exposure, no such enrichment for development-related genes could be detected (Supplementary Figure 4B and D). This suggests that the changes in the methylation level of these genes might serve another purpose than an immediate adjustment of their transcriptional activity. Alternatively, the role of CLF in the cold stress response may be masked by its paralogue SWN, as both proteins have overlapping functions, at least for developmental processes (Chanvivattana *et al*., 2004).

Similarly, while many cold-inducible genes underwent a gain of H3K4me3 upon cold exposure, this could not be generalized to all up-regulated genes. This was also observed for other abiotic stresses (Sani *et al*., 2013; Yamaguchi *et al*., 2021). The correlation analyses between differential methylation and expression suggest that both phenomena have different dynamics, with expression changes occurring prior to methylation status alterations. This would indicate that H3K4me3 gain is not necessary for the initiation of the transcriptional activation but it might positively feed back into it, as genes gaining H3K4me3 displayed a higher magnitude of induction than non DM genes (Figure 4C). While H3K4me3 has long been described as being necessary for transcription initiation, this idea has recently been refuted (Shilatifard, 2012; Lauberth *et al*., 2013; Wang *et al*., 2023). Instead, H3K4me3 was demonstrated to prevent RNA polymerase II pausing, thereby accelerating elongation. This suggests that higher H3K4me3 levels would lead to higher accumulation of transcripts, as observed in this study for genes gaining H3K4me3 upon cold exposure (Figure 4B).

Despite the observed correlations between differential methylation and differential expression, it is important to note that numerous cold-regulated genes did not undergo differential methylation and vice-versa (Supplementary Figure 5), indicating that differential methylation is neither required for differential expression nor it’s a direct consequence. However, differential methylation might allow for a larger magnitude of induction of cold-responsive genes, as both H3K4me3-gaining and H3K27me3-losing genes displayed slightly higher fold-change of gene expression than non-differentially methylated genes (Figures 4 and 5). Alternatively, the limited overlap between differential methylation and expression might be explained by the fact that all the epigenomic and transcriptomic experiments of the current study have been performed on whole seedlings. This prevents us from testing whether different tissues or cell types respond differently to lower temperatures. In tomatoes for example, nitrogen treatment triggered H3K4me3 and H3K27me3 differential methylation on distinct sets of genes in shoots and roots (Julian, Patrick and Li, 2023). Performing similar investigations in a tissue-specific approach might allow us to decipher more precisely the relationship between histone methylation and transcriptional activity.

Uncovering the exact potential role of differential histone methylation in the response to cold will require the identification of the mechanisms controlling it. For both H3K4me3 and H3K27me3, differential methylation was not associated with altered nucleosome density (data not shown), suggesting that differential methylation is due to active mechanisms rather than H3 depletion or accumulation. H3K4me3 is deposited by methyltransferases, which are known to act redundantly in *Arabidopsis thaliana* (Chen *et al*., 2017; Cheng *et al*., 2020). According to the transcriptomic data generated in this study and previously generated data, both *ATX1* and *ATX4* are induced by exposure to low temperatures (Supplementary Figure 6A, Vyse et *al.* 2020), suggesting them as first candidates. In particular, ATX1 has already been shown to deposit H3K4me3 on specific genes upon cold treatment (Miura, Renhu and Suzaki, 2020). H3K27me3 loss upon heat has been demonstrated to be redundantly controlled by JMJ30, JMJ32, ELF6 and REF6 (Yamaguchi *et al*., 2021), the same methyltransferases might therefore regulate H3K27me3 levels during cold stress. In particular, both ELF6, JMJ13 and JMJ30 were found to be induced during cold exposure (Supplementary Figure 6B, Vyse et *al.* 2020). It would therefore be of high interest to examine the cold tolerance abilities and transcriptional response of such mutants to cold exposure.

In conclusion, this study provides a genome wide perspective on cold-triggered histone methylation dynamics and demonstrates that H3K4me3 and H3K27me3 differential methylations are independent from one another. H3K4me3 correlates more strongly with differential expression and appears to regulate immediate stress responses, while H3K27me3 might contribute to longer term responses such as developmental adaptation. As reduced H3K27me3 levels did not impact the transcriptional activity of cold-responsive genes, further work is required to finally elucidate the role played by this repressive mark at those genes. It would especially interesting to examine whether it might contribute to deacclimation processes.

## Abbreviations

COR: Cold Responsive
DE: Differentially Expressed
DM: Differentially Methylated
GO: Gene Ontology
H3K4me3: Histone 3 Lysine 4 trimethylation
H3K27me3: Histone 3 Lysine 27 trimethylation
TES: Transcription End Site
TSS: Transcription Start Site

## 6 Author Contributions

LF and DS conceived the project. LF performed most of the experiments, analyzed and interpreted the data. NFK performed the western blots experiments. ABK performed the electrolyte leakage experiments. XX helped to prepare the ChIP-seq libraries. KK contributed to the ChIP-seq design and analysis. LF and DS drafted the manuscript, which was then revised by all authors.

## 8 Acknowledgments

The authors would like to thank Jose M. Muino for his precious expertise in ChIP-seq analysis and the HPC Service of ZEDAT, Freie Universität Berlin, for computing time.

## 11 Data Availability Statement

The datasets generated for this study has been deposited in NCBI’s Gene Expression Omnibus and are accessible through GEO Series accession number GSE255445 (https://www.ncbi.nlm.nih.gov/geo/query/acc.cgi?&acc=GSE255445)

