## Supplementary Figures for "Cold stress induces a rapid redistribution of the antagonistic marks H3K4me3 and H3K27me3 in *Arabidopsis thaliana*"

Supplementary Material

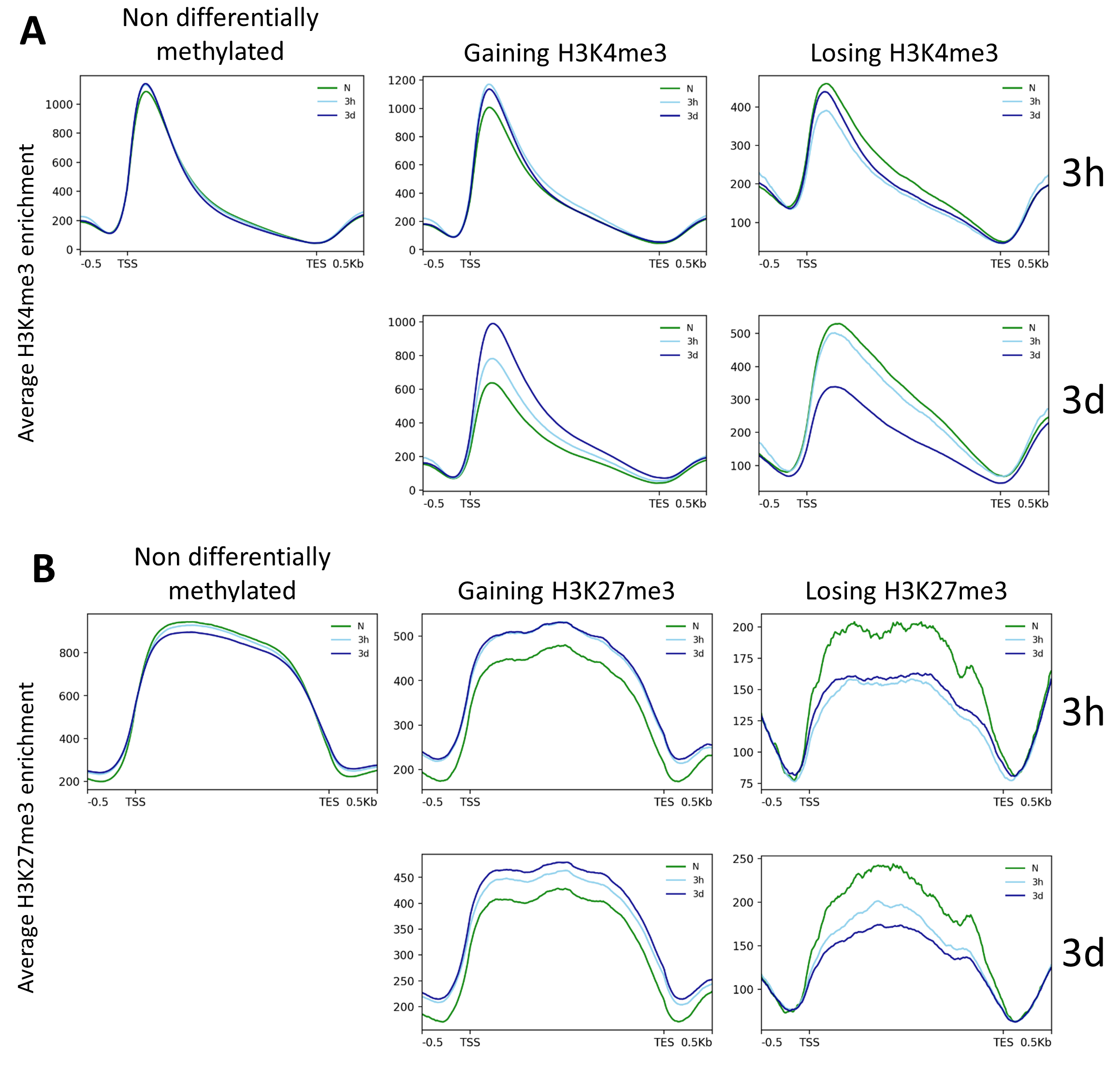

**Sup. Figure 1: Methylation levels on differentially methylated genes.** Metagene plots showing the levels of H3K4me3 **(A)** and H3K27me3 **(B)** on non-differentially methylated genes (left panel), genes gaining (middle panel) or losing the respective mark (right panel), after three hours (3h) or three days (3d) of cold exposure. Differentially methylated genes are genes showing an absolute log2 fold change >= 0.5 of the respective methylation mark on a region spanning from the TSS to the TES.

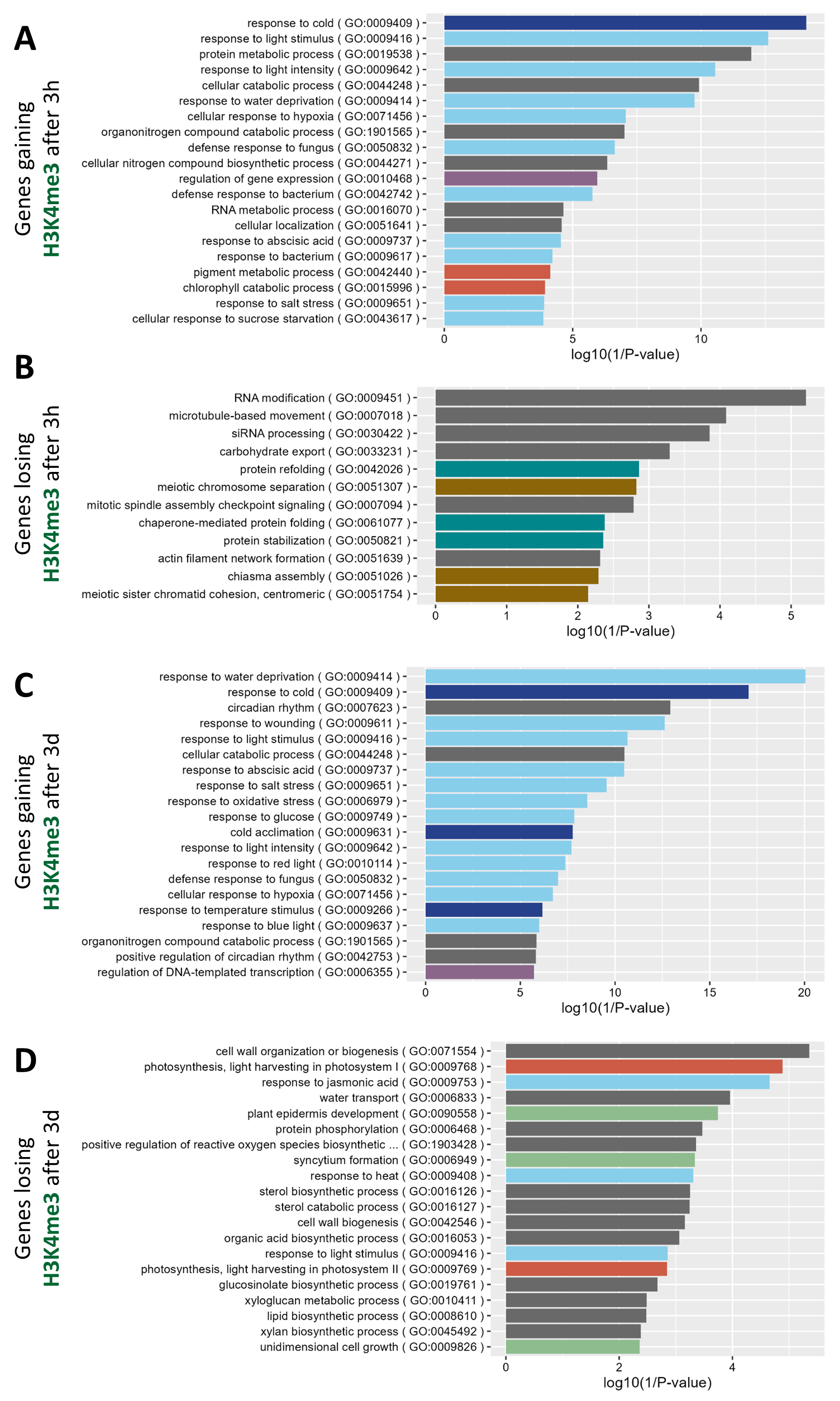

**Sup. Figure 2: Gene ontology (GO) analysis of genes undergoing H3K4me3 differential methylation upon cold exposure.** Enriched biological process GO terms of genes gaining (**(A)** and **(C)**) or losing (**(B)** and **(D)**) H3K4me3 after three hours (**(A)** and **(B)**) or three days (**(C)** and **(D)**) of cold exposure. Colors indicate the broad category of the term: light blue: stress response, dark blue: temperature response, purple: regulation of transcription, orange: photosynthesis, turquoise: protein folding, gold: chromatin, green: development.

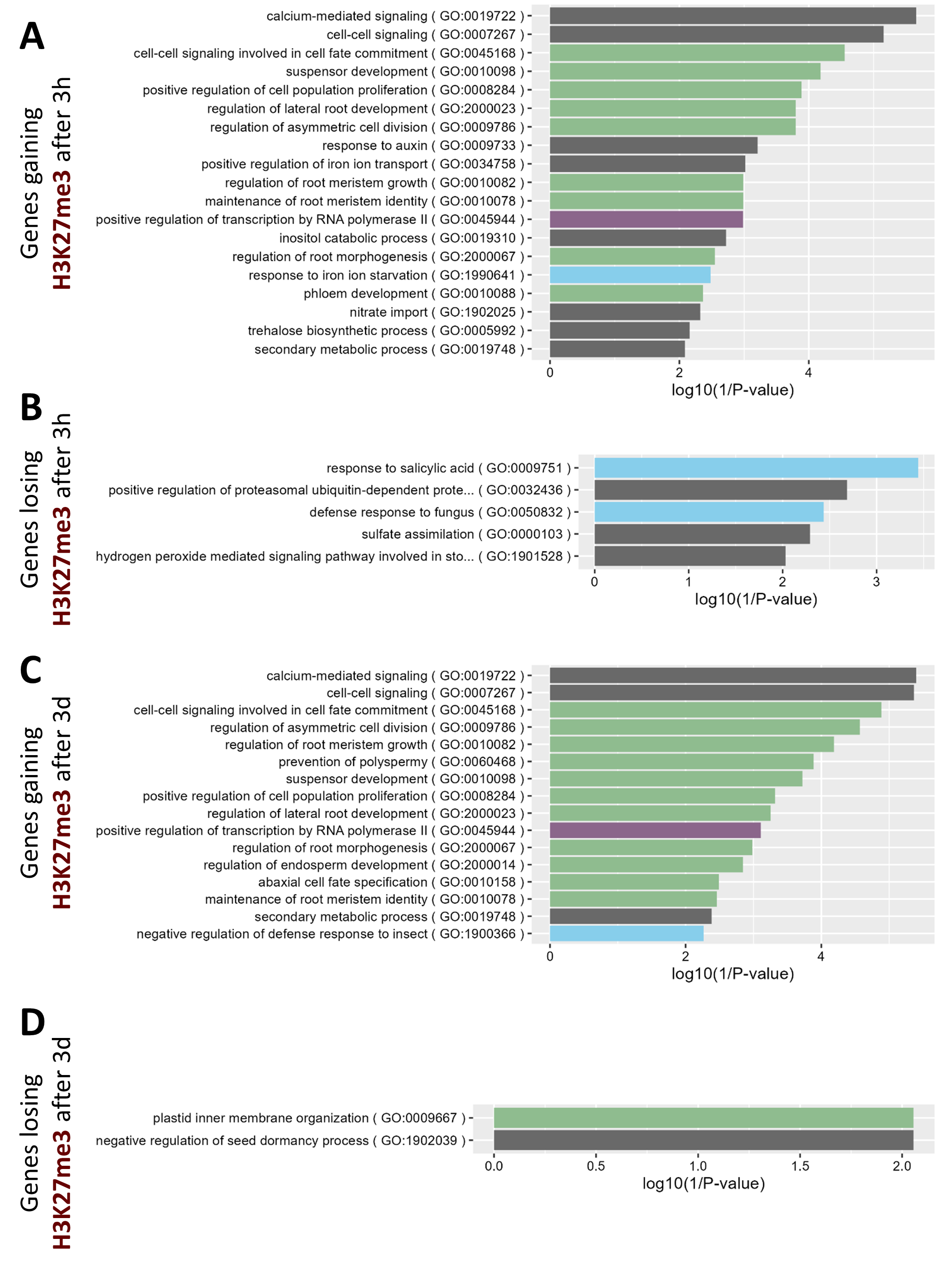

**Sup. Figure 3: Gene ontology (GO) analysis of genes undergoing H3K27me3 differential methylation upon cold exposure.** Enriched biological process GO terms of genes gaining (**(A)** and **(C)**) or losing (**(B)** and **(D)**) H3K27me3 after three hours (**(A)** and **(B)**) or three days (**(C)** and **(D)**) of cold exposure. Colors indicate the broad category of the term: light blue: stress response, purple: regulation of transcription, green: development.

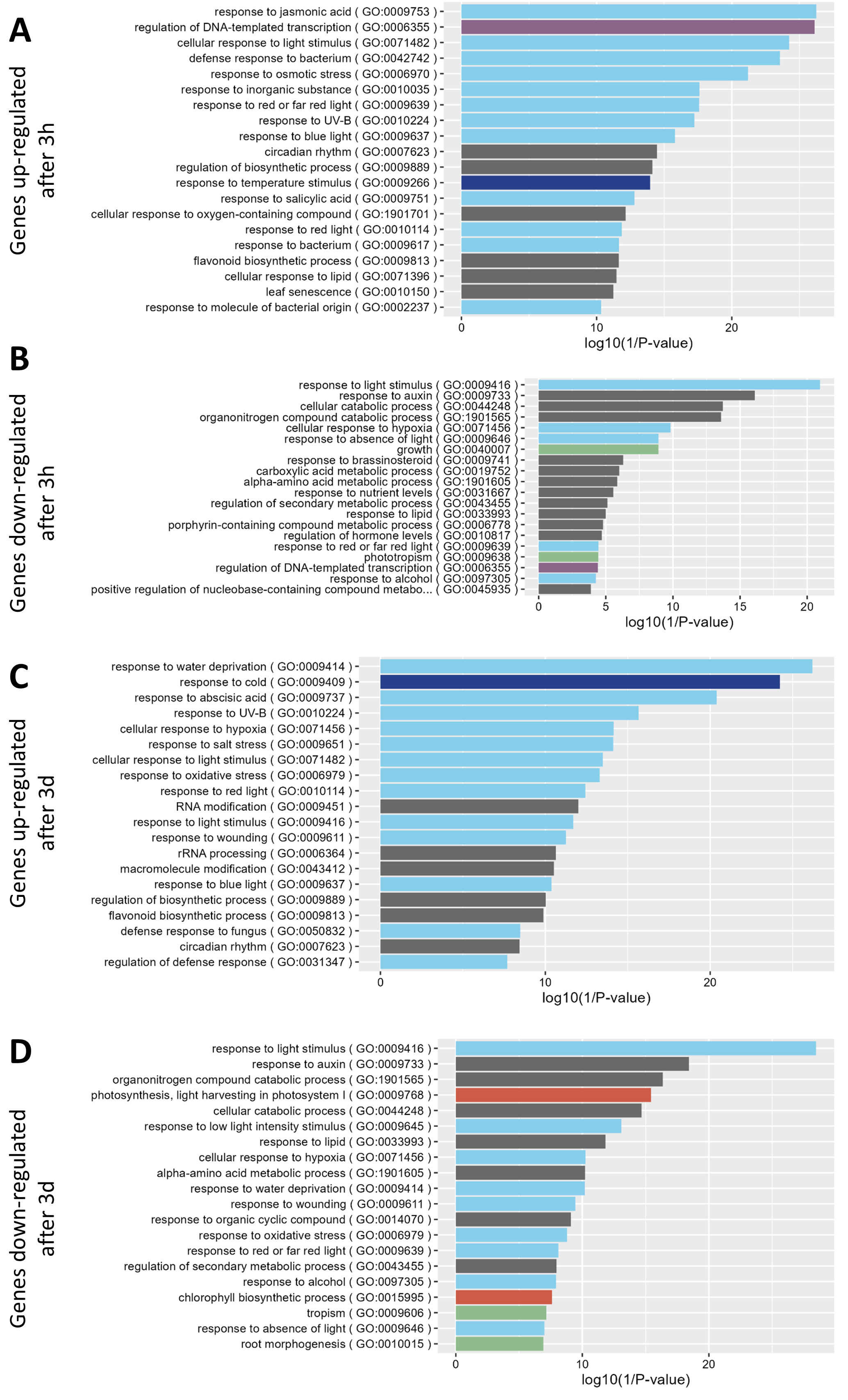

**Sup. Figure 4: Gene ontology (GO) analysis of differentially expressed genes upon cold exposure.** Enriched biological process GO terms of up- (**(A)** and **(C)**) or down-regulated genes (**(B)** and **(D)**) after three hours (**(A)** and **(B)**) or three days (**(C)** and **(D)**) of cold exposure. Colors indicate the broad category of the term: light blue: stress response, dark blue: temperature response, purple: regulation of transcription, orange: photosynthesis, green: development.

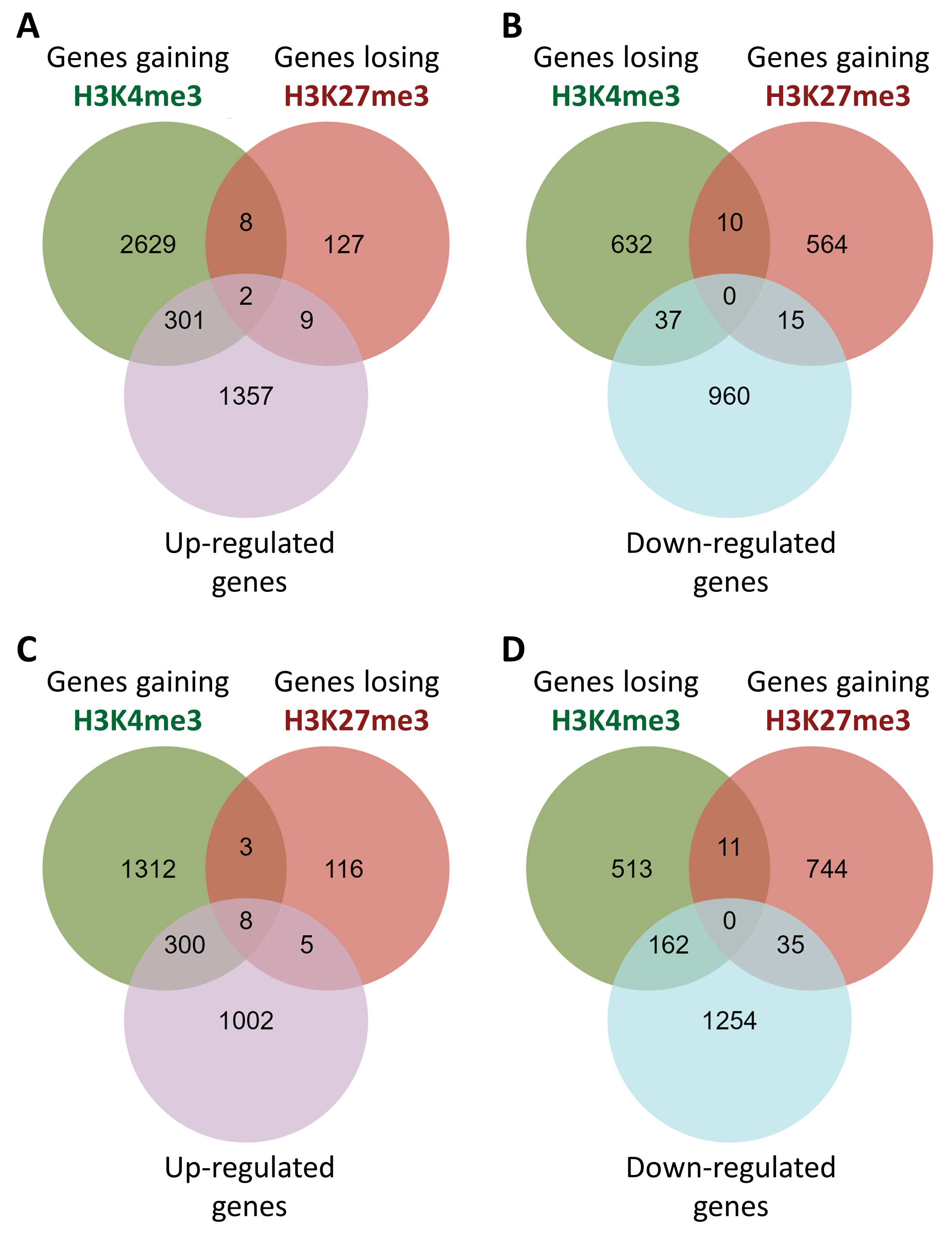

**Sup. Figure 5: Differentially methylated and differentially expressed genes partially onverlap.** Venn diagrams showing the overlaps between up-regulated genes, genes gaining H3K4me3 or losing H3K27me3 after 3h **(A)** or 3d **(C)** of cold exposure. Venn diagrams showing the overlaps between down-regulated genes, genes losing H3K4me3 or gaining H3K27me3 after 3h **(B)** or 3d **(D)** of cold exposure.

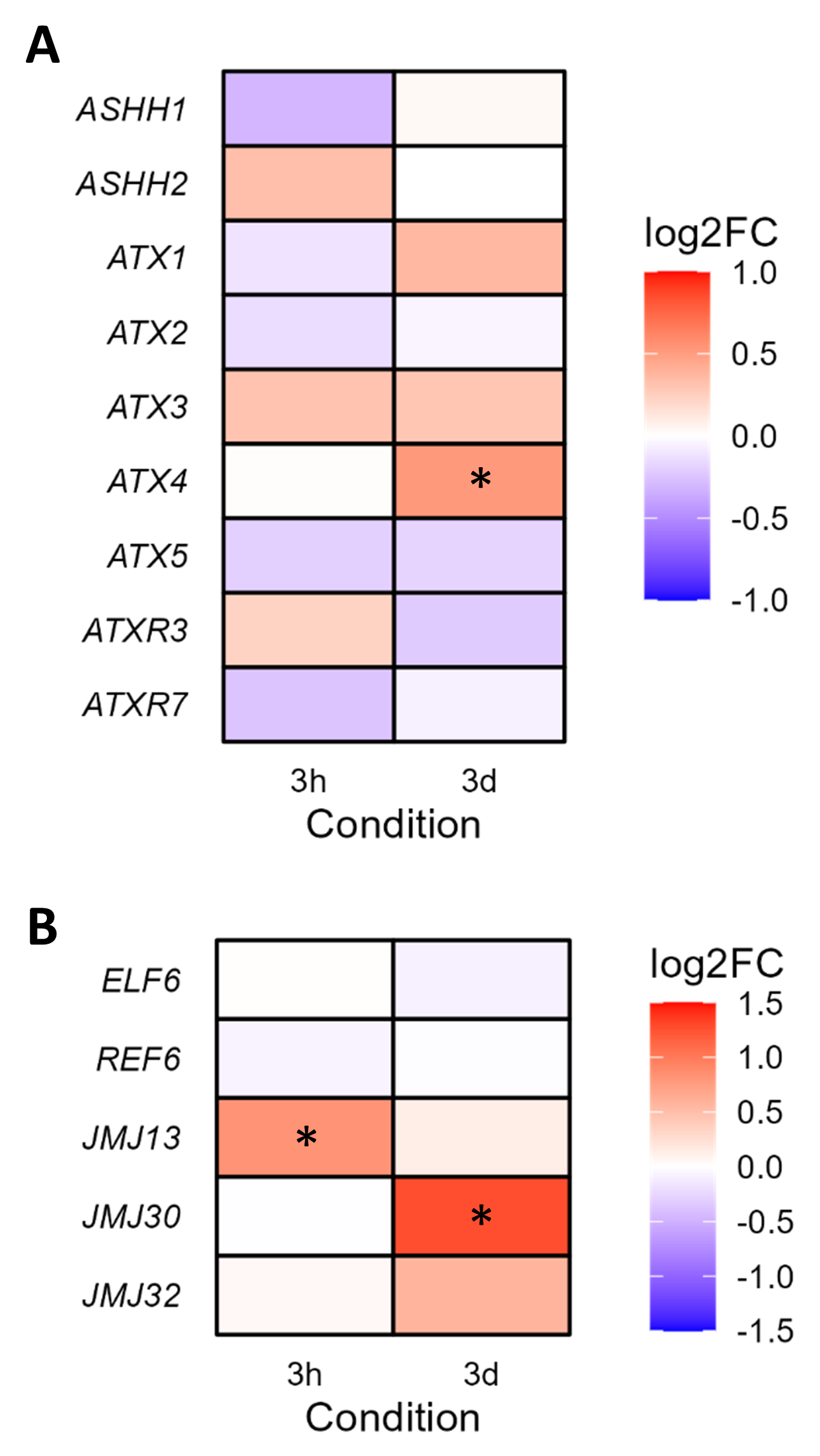

**Sup. Figure 6: Expression changes of genes encoding H3K4me3 methylases (A) or H3K27me3 demethylases (B).** Plants were exposed to cold for either 3h or 3d and the gene expression was analyzed using a RNA-seq. The color indicates the log2 fold change compared to non-treated condition. * indicates p-adj < 0.05
